# Whole genome sequencing provides evidence of two biologically and clinically distinct entities of asymptomatic monoclonal gammopathies: progressive versus stable myeloma precursor condition

**DOI:** 10.1101/2020.11.06.372011

**Authors:** Bénedith Oben, Guy Froyen, Kylee H. Maclachlan, Daniel Leongamornlert, Federico Abascal, Binbin Zheng-Lin, Venkata Yellapantula, Andriy Derkach, Ellen Geerdens, Benjamin T. Diamond, Ingrid Arijs, Brigitte Maes, Kimberly Vanhees, Malin Hultcrantz, Elisabet E. Manasanch, Dickran Kazandjian, Ahmet Dogan, Yanming Zhang, Aneta Mikulasova, Brian Walker, Gareth Morgan, Peter J. Campbell, Ola Landgren, Jean-Luc Rummens, Niccolò Bolli, Francesco Maura

## Abstract

Multiple myeloma (MM) is consistently preceded by precursor conditions recognized clinically as monoclonal gammopathy of undetermined significance (MGUS) or smoldering myeloma (SMM). We interrogate, for the first time, the whole genome sequence (WGS) profile of 18 MGUS and compare them with those from 14 SMMs and 80 MMs. We show that cases with a non-progressing, clinically stable myeloma precursor condition (n=15) are characterized by later initiation in the patient’s life and by the absence of myeloma defining genomic events including: chromothripsis, templated insertions, mutations in driver genes, aneuploidy, and canonical APOBEC mutational activity. This data provides evidence that WGS can be used to recognize two biologically and clinically distinct myeloma precursor entities that are either progressive or stable.

## Introduction

Multiple myeloma (MM) is the second most common hematological malignancy and is consistently preceded by the asymptomatic expansion of clonal plasma cells, termed either monoclonal gammopathy of undetermined significance (MGUS) or smoldering myeloma (SMM).^1-6^ These two precursor conditions are found in 2-3% of the general population aged older than 40 years. Only a small fraction of MGUS will ultimately progress to MM, whereas approximately 65% of persons with SMM will progress within 10 years of initial diagnosis.^2,4^ Currently, the differentiation between MGUS and SMM is based on indirect measures and surrogate markers of disease burden.^5,6^ While these features are reasonably accurate in defining a SMM high-risk disease group and its average risk of progression,^7^ they perform significantly less well in predicting risk for the group of patients with low disease burden (e.g. intermediate- and low-risk SMM). Moreover, they do not provide a personalized assessment of risk for the individual patient.^8-10^

In the last decades, next generation sequencing (NGS) approaches have facilitated major progress in deciphering the genomic complexity of MM and its precursor conditions.^11,12^ Mutations in driver genes and structural events (e.g. *MYC* translocations) have been reported to be infrequent in precursor conditions compared to MM, and their presence has been suggested to confer a higher risk of progression.^5,13-20^ However, these studies had two major limitations: 1) they were based on exome/targeted sequencing approaches and hence were not able to fully capture the landscape of myeloma defining genomic events; 2) they focused mostly on SMM and did not include MGUS.

Whole genome sequencing (WGS) has emerged as the most comprehensive approach to characterize MM and myeloma precursor conditions due to its ability to interrogate the full repertoire of myeloma defining genomic events including: single nucleotide variants (SNVs), mutational signatures, copy number variants (CNVs), and structural variants (SVs).^9,13,21-25^ However, the use of WGS on myeloma precursor conditions has been historically limited by the low clonal bone marrow plasma cell (BMPC) percentage, and therefore the availability of tumor DNA. To circumvent this challenge, in MGUS and SMM samples with low cellularity included in this study, we applied multi-parameter flow-cytometry sorting and an innovative low input WGS approach (**Figure 1A**) able to characterize the genomic landscape of normal tissue from a few thousand cells.^26,27^ Thanks to this novel methodology, we have been able to perform the first comprehensive comparison at the whole genome level between low and intermediate risk myeloma precursor condition and MM. The results of the study provide strong evidence for two biologically and clinically distinct myeloma precursor condition entities: (1) a progressive myeloma precursor condition, which is a clonal entity in which myeloma defining genomic events have already been acquired at the time of sampling and associated with an extremely high risk of progression to MM; and (2) a stable myeloma precursor condition, in which myeloma defining genomic events are rare and that follows an indolent clinical course.

**Figure 1.**
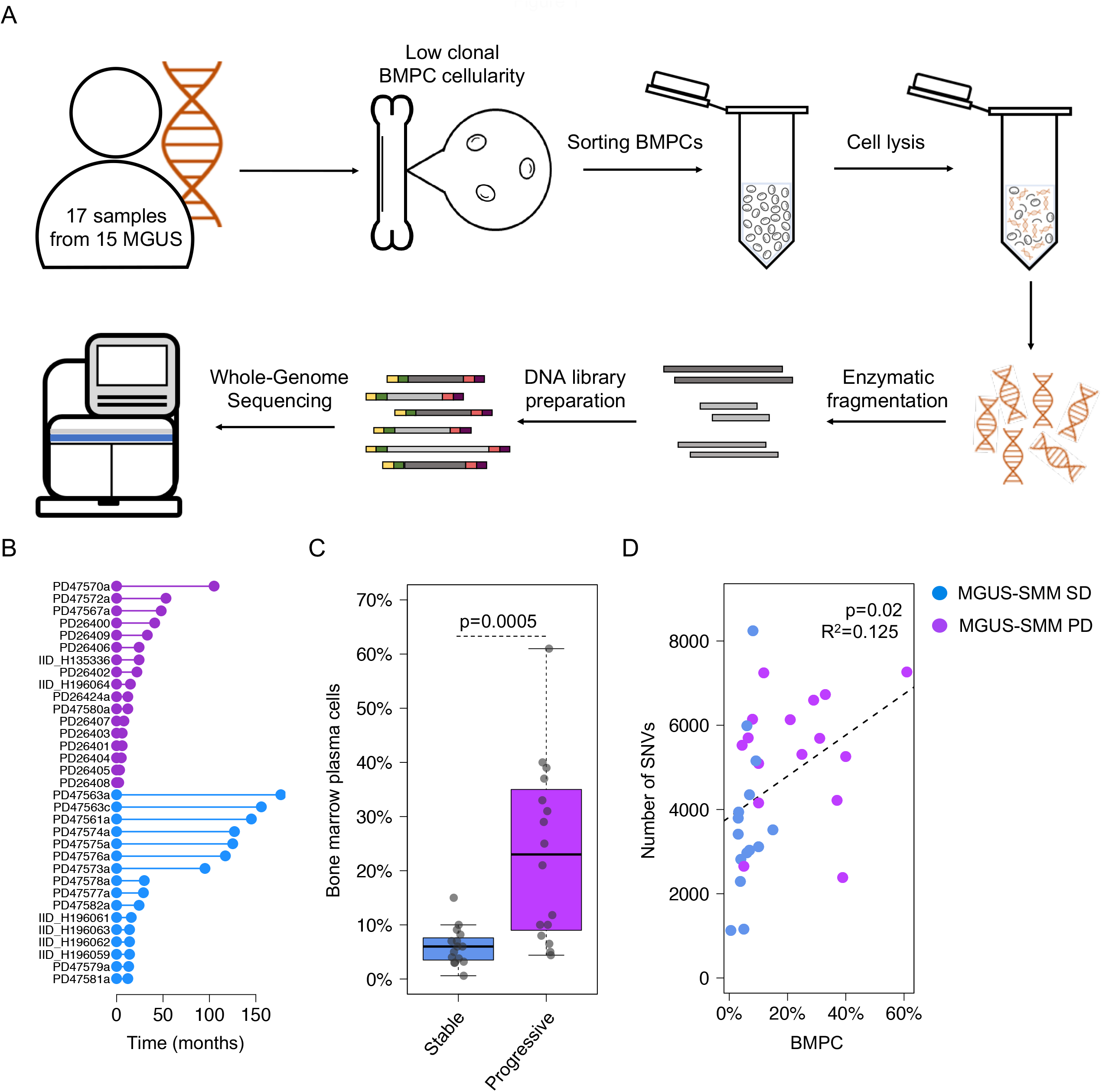
Summary of all patients with myeloma precursor condition included in the study. A) A cartoon summarizing the low input WGS approach. B) Follow up time for all patients with myeloma precursor condition included in the study. Purple and blue lines and dots reflect patients that progressed to MM and hadn’t had progression at the time of study, respectively. C) Comparison of bone marrow plasma cell infiltration between stable and progressive myeloma precursor condition. p value was generated using Wilcoxon rank-sum test. D) Correlation between bone marrow plasma cell (BMPC) infiltration and mutational burden in myeloma precursor condition. p value and R^2^ were estimated using *lm* R function (linear regression).

## Results

### Single nucleotide-based substitution mutational signatures

We interrogated the WGS profile of 32 patients with myeloma precursor condition defined according to the International Myeloma Working Group 2014 criteria (MGUS=18; SMM=14).^7^ Only one of the 14 SMM patients was defined as high-risk based on the Mayo Clinic prognostic model (PD26424a).^3^ None of the patients showed signs of progression at the time of sample collection and none of the SMM cases had a bone lesion on either skeletal radiography, CT, or PET-CT.^7^ After a median follow up of 24 months from sample collection (range: 2-177), 17 out of 32 (53%) patients with a myeloma precursor condition progressed to MM and started anti-MM treatment [13/14 SMM and 4/18 defined clinically as MGUS; median time to progression: 14.5 months (range: 2-105)] (**Figure 1B; Supplemental Tables 1**-**3**). In the current study, patients who had subsequent progression to MM are defined as having “progressive myeloma precursor condition”. Patients with prolonged clinical stability (at least 1 year of follow up without progression to MM) were defined as “stable myeloma precursor condition” [mean follow up 69 months (range: 12-177)]. The stable myeloma precursors had a significantly lower BMPC infiltration compared to progressive cases (Wilcoxon rank-sum test p=0.0005; **Figure 1C**), likely reflecting the higher proportion of MGUS (Fisher’s exact test p<0.0001). Stable myeloma precursor disease had a median mutational burden of 3406 SNVs (range 1130-8244), that is significantly lower in comparison with the progressive myeloma precursor disease (5518; range 2385-7257; Wilcoxon rank-sum test p=0.034) and MM (5482 range 982-15738; Wilcoxon rank-sum test p=0.005). The mutational burden was weakly correlated with the BMPC infiltration(p=0.02 and R^2^=0.125; **Figure 1D**). To interrogate if the difference in overall mutational burden reflects the activity of different mutational processes, we explored the single-base substitution (SBS) mutational signature landscape of stable myeloma precursor condition in comparison to that of progressive myeloma precursor condition and MM.^28,29^ Running *de novo* signature extraction across the entire cohort of plasma cell disorders (n=112), all main MM mutational signatures were identified: aging (SBS1 and SBS5), AID (SBS9), SBS8, SBS18, and APOBEC (SBS2 and SBS13) (**Supplemental Figure 1** and **Supplemental Table 4**).^13,25,29^ APOBEC emerged as the most differentially active mutational process across the three groups (**Figure 2A**). Interestingly, only 13% (2/15) of stable myeloma precursor condition cases showed significant evidence of APOBEC activity, in comparison with 82% (14/17) and 85% (68/80) of patients with progressive myeloma precursor condition (p=0.009) and MM (p=0.0006), respectively (**Figure 2A-B**). Furthermore, the two stable cases with a detectable APOBEC signature were characterized by a high APOBEC3A:3B ratio ^25,28,30^ a feature which defines a group of MAF-translocated MM patients characterized by intense and early APOBEC activity ^25,31,32^. The mutational activity pattern in this group is clearly different from that observed in the majority of progressive myeloma precursors and MM cases, which are characterized by a ~1:1 APOBEC3A:3B ratio (**Figure 2C**). In line with this mutational profile, both stable myeloma precursor cases with APOBEC3A:3B ratio had a translocation between *IGH* and *MAFB* reinforcing the notion that APOBEC activity must be assessed in the light of the APOBEC3A:3B ratio, as this appear to highlight different biological and clinical disease entities.

**Figure 2.**
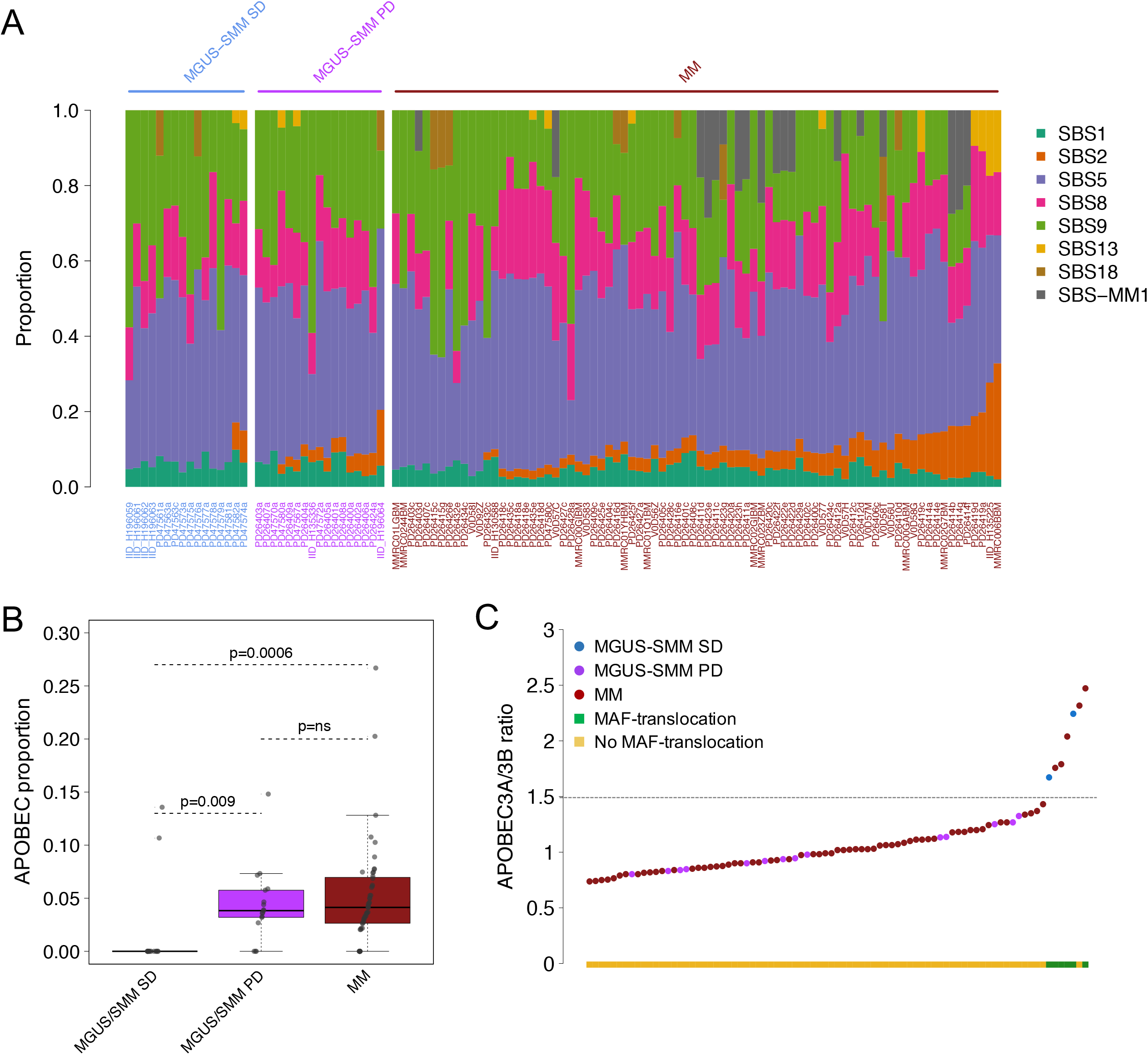
Mutational signature landscape of multiple myeloma (MM) and myeloma precursor condition. A) Mutational signature contribution across all WGS samples included in the study. B) Comparison of APOBEC mutational contribution (SBS2 + SBS13) between MM, progressive and stable myeloma precursors. p values were calculated using Wilcoxon rank-sum test. C) APOBEC3A:3B ratio of all patients included in the study having detectable APOBEC activity. Blue, purple and brown dots represent stable, progressive myeloma precursors and MM, respectively. The green and yellow boxes on the x-axis reflect cases with and without translocations involving *MAF*/*MAFB*, respectively. MGUS: monoclonal gammopathy of undetermined significance; SMM: smoldering multiple myeloma; SD: stable disease; PD: progressive disease.

### Single nucleotide variants in known myeloma driver genes

By integrating indels and SNVs, we explored the distribution of mutations in 80 known myeloma driver genes in the myeloma precursor conditions (**Supplemental Table 5**).^12,33^ To increase the power of the investigation, we included two different whole exome sequencing (WXS) datasets: the first comprised of 33 MGUS (2 of which progressed) and the second comprised of 947 newly diagnosed MM enrolled in the CoMMpass trial (version IA13, Multiple Myeloma Research Foundation Personalized Medicine Initiative).^19,33^ Overall, analogous to previous reports, patients with stable myeloma precursor condition were characterized by a significantly lower number of mutations in known myeloma driver genes compared with progressive myeloma precursor condition (Wilcoxon rank-sum test p=0.002) and MM (Wilcoxon rank-sum test p<0.0001) (**Figure 3A-B**).^14,17,19^ Investigating the patterns of positive selection across the different stages of disease using *dNdScv*,^34^ we observed a significant signal indicative of positive selection in the known myeloma driver genes in progressive myeloma precursor condition and MM, but this pattern was not seen in the stable myeloma precursor condition (**Supplemental Data 1**). Calculating the per-gene confidence intervals for dN/dS values under the *dNdScv* model using profile likelihood, only mutations in *HIST1H1E* (n=2) showed evidence for positive selection among the stable myeloma precursor conditions. The same analysis among the progressive myeloma precursor conditions and MM (both WGS and WXS) showed that multiple driver genes were under selective pressure, including mutations involving MAPK and NFkB pathways, and tumor suppressor genes such as *TP53* (**Supplemental Data 1**).

**Figure 3.**
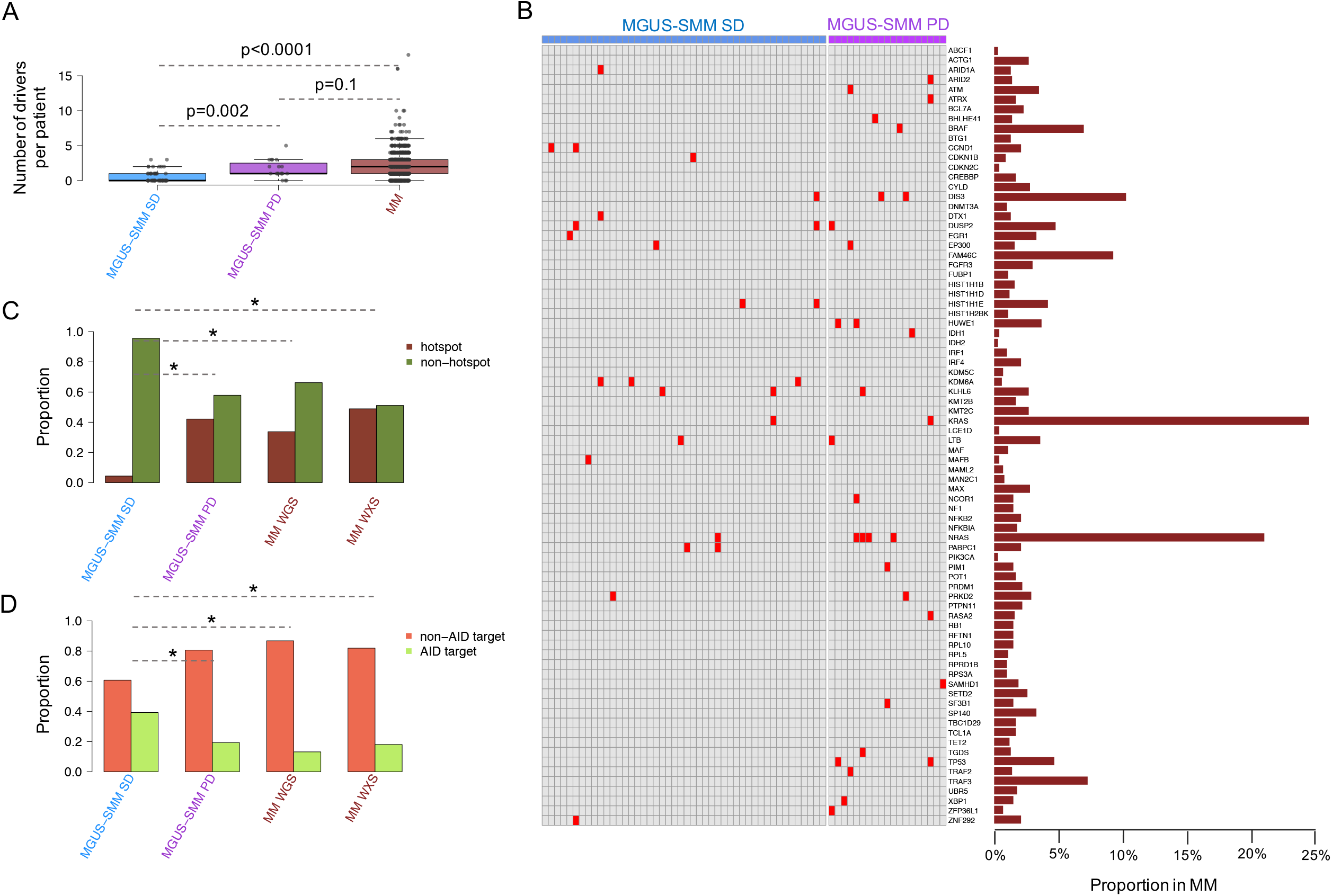
Mutations in myeloma driver genes. A-B) Prevalence and distribution of nonsynonymous mutations in driver genes (n=80) across stable (blue) and progressive (purple) myeloma precursor condition and multiple myeloma (brown). C) Proportion of cases with at least one significant known hotspot mutation within myeloma driver genes in stable and progressive myeloma precursor condition and MM. D) Proportion of mutations in driver genes involving known AID targets in stable and progressive myeloma precursor condition and MM. SD: stable disease, PD: progressive disease. Asterisks in C and D indicate a p<0.01 under Fisher’s exact test of proportions.

To further characterize the mutational driver landscape of myeloma precursor conditions we ran *sitednds* to identify known mutational hotspots within known myeloma driver genes. In line with the previous analysis, patients with stable myeloma precursor condition were characterized by a lower number of mutations in known driver hotspots compared to progressive myeloma precursor condition and MM (**Figure 3C**; **Supplemental Data 1**). Finally, in line with their mutational signature profile and with the absence of APOBEC activity, stable myeloma precursor condition had a higher proportion of mutations within known AID targets compared to progressive myeloma precursor condition and MM (**Figure 3D**).^28,33,35^ Overall, these results suggest that the mutational landscape of stable myeloma precursor condition is significantly different in comparison to progressive myeloma precursor condition and MM, in terms of both number of mutations in myeloma driver genes and the mutational processes involved.

### Copy number changes and structural variants

When exploring the cytogenetic landscape, no significant differences in recurrent aneuploidies were found between the progressive myeloma precursor condition and the MM cases. In comparison to progressor condition and to MM, patients with stable myeloma precursor condition were characterized by a significantly lower prevalence of known recurrent MM chromosomal abnormalities including gain1q, del6q, del8p, gain8q24 and del16q; **Supplemental Figure 2** and **Supplemental Table 6**). This observation was validated combining our WGS cohort with additional SNP array copy number data from 66 stable myeloma precursor condition, 2 progressive myeloma precursor condition, and 148 MM patients, respectively (p<0.001 for these recurrent abnormalities; **Figure 4** and **Supplemental Table 7-8**).

**Figure 4.**
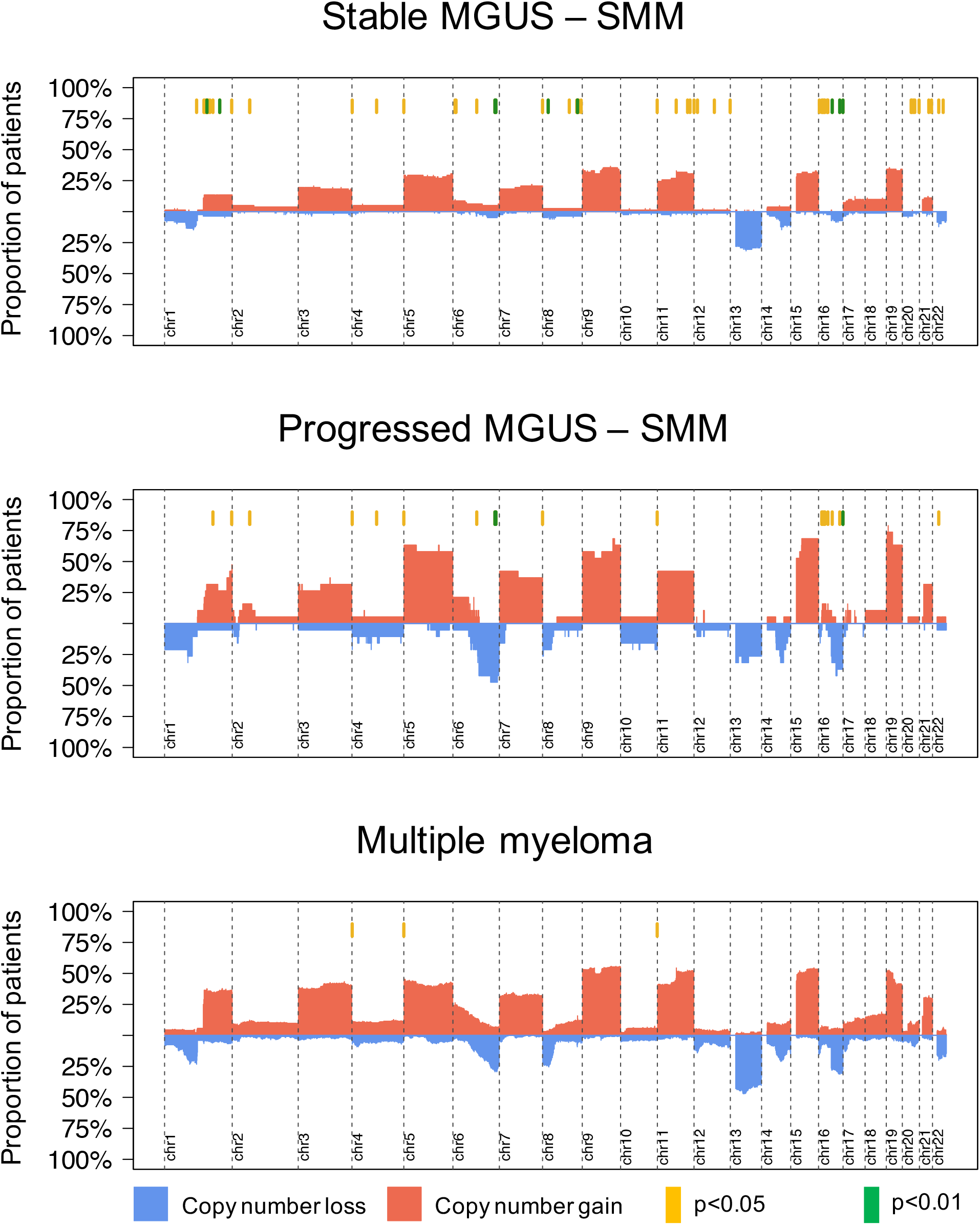
Copy number profile of myeloma precursor condition and multiple myeloma (MM). Cumulative copy number profile of all patients with either WGS or SNP array data available. Cases were grouped according to their clinical stage: stable and progressive myeloma precursor condition and MM. Red and blue bars reflect chromosomal gain and loss, respectively. Yellow and green lines on the top of each graph represent GISTIC peaks with a significantly different prevalence across the three stages (yellow: Fisher’s exact test p<0.05 and green: q<0.1). On the first, second and third cumulative plots we reported the significant difference between: stable myeloma precursor condition *vs* MM, stable myeloma precursor condition *vs* progressive myeloma precursor condition and progressive myeloma precursor condition *vs* MM, respectively.

To further characterize differences in myeloma defining genomic events between stable versus progressive myeloma precursor condition and MM, we leveraged the comprehensive resolution of WGS to explore the distribution and prevalence of SVs and complex SV events, known to play a critical role in MM pathogenesis. Stable myeloma precursor cases were characterized by a lower SV burden overall. This was true for single SVs (Wilcoxon rank-sum test p=0.0005), but was even more striking for complex SVs (Wilcoxon rank-sum test p<0.0001; **Figure 5A**).^22,24,36,37^ Only one stable myeloma precursor case had a chromothripsis event, and none had evidence of templated insertions between either two, or more than two chromosomes. This scenario was significantly different in progressive myeloma precursor condition, where chromothripsis and templated insertions were detected in 8/17 (47%; p<0.001) and 7/17 (41%; p<0.001), respectively. Overall, the progressive myeloma precursor condition SV landscape was similar to that observed in MM, itself (**Figure 5B-C**). This finding was confirmed by looking at the genomic distribution of SV: in progressive precursors, and to a greater extent in MM, the distribution was significantly associated with H3K27a and chromatin accessibility loci (**Supplemental Figure 3**).^24^

**Figure 5.**
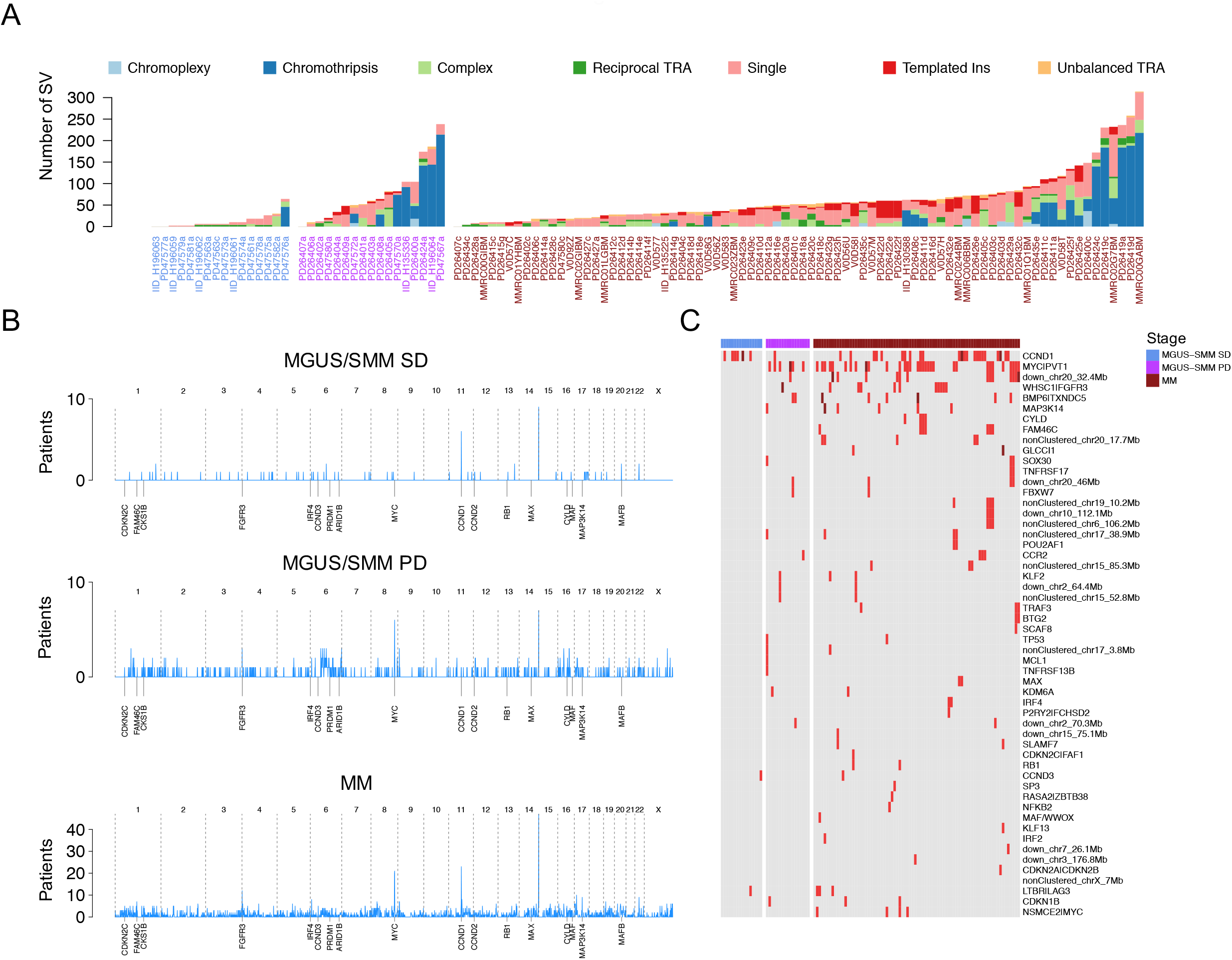
Landscape of structural variants (SV) in multiple myeloma (MM) and myeloma precursor condition. A) Prevalence of single and complex SV events across all cases included in this study. Blue, purple and brown x-axis labels represent stable, progressive myeloma precursors and MM respectively. Ins: insertion, TRA: translocation. B) Genome-wide density of SV breakpoints across stable, progressive myeloma precursors and MM. Each patient genome was divided in bins of 1 Mb and, in case of presence of multiple SV breakpoints, only one breakpoint was counted. C) Prevalence of 69 known SV hotspots across stable and progressive myeloma precursors, and MM. SD; stable disease, PD; progressive disease.

We analyzed hotspots hit by recurrent SV in our case series. Sixty-nine hotspots were identified in 752 low-coverage long-insert WGS cases from the CoMMpass data set.^21,24^ The median number of these SV hotspots per patient was significantly lower among stable myeloma precursor condition compared to MM (Wilcoxon rank-sum test p=0.0001; **Supplemental Figure 4**). Among the stable myeloma precursor condition cases, we identified only 11 SV hotspots: all translocations between the *IGH* locus and C*CND1* (n=7), *MAFB* (n=2), *CCND3* (n=1), and *LTBR*|*LAG3* (n=1). Of note, none of the stable myeloma precursor condition cases had any SVs involving the *MYC*/*PVT1* hotspot^13,19^ in sharp contrast with 35% (6/17) in progressive precursor condition cases and 32/80 (40%) MM (Fisher’s exact test p=0.03 and p=0.003, respectively). Overall, progressive myeloma precursor condition did not show any significant differences in SV hotspot prevalence compared to either MM or stable myeloma precursor condition.

### Time lag between initiation and sample collection

Considering myeloma defining genomic events (i.e. SNVs, CNVs, SVs and mutational signatures), stable myeloma precursor condition emerged as a distinct genomic entity compared to MM. In contrast, the progressive myeloma precursor condition demonstrated a genomic profile extremely similar to that of MM. This absence of myeloma defining genomic events among stable cases could be due to two possible explanations. Firstly, the early detection of the clone by serum protein electrophoresis and consequent earlier sample collection in the course of disease might have introduced a temporal bias into our analysis (i.e. the earlier the plasma cell clonal detection, the lower its genomic complexity). Alternatively, stable cases represent a distinct biological entity, characterized by few genomic aberrations and a low propensity to acquire additional abnormalities associated with progression. To identify the most likely model, we leveraged the molecular-clock approach, recently developed to time landmark events in both cancers and normal tissues.^25-27,38,39^ Notably, this approach is based on the SBS1 and SBS5 mutational burden pre- and post-chromosomal gain to estimate the time lag between cancer-initiating gains and sample collection. Previous MM molecular time estimates^25^ are in line with a long lag time from initiation to development.^40,41^ For this study, this analysis was performed in three main steps. Firstly, we used the Dirichlet process-derived clonal mutational burden of each patient to estimate the relative time of acquisition of each large chromosomal gain. In this way, we could identify large chromosomal gains occurring within the same time window. Then, we estimated the contribution of each mutational signature; collapsing together duplicated and non-duplicated mutations within the earliest multi-chromosomal gain event in each patient. Finally, we estimated the SBS1- and SBS5-based molecular time of each early multi-gain event and converted it to patient years. Overall the age at sampling was not significantly different between MM, stable and progressive myeloma precursor condition (**Figure 6A**). However, when we used the molecular timing approach we were able to show that the stable myeloma precursor condition cases had a significantly different temporal pattern, in which multi-gain events occurred later in the patient’s life (median 53.5 years; range 28-65) compared to the progressive myeloma precursor condition cases (median 28 years range 5-46) and MM cases (median 20.5; range 9-56) (**Figure 6B-C**). These data argue against a temporal bias created by early sample collection relative to disease initiation in non-progressing samples. Instead the results suggest that while these stable entities may eventually progress to MM, based on these temporal estimates, this would be predicted to occur at average ages of 90-100 years of age. Overall, our temporal estimates suggest that stable myeloma precursor condition represents a different biological entity; one that is acquired at a later age in life, without myeloma defining genomic events, and with a much lower tendency to progress compared to progressive myeloma precursor condition.

**Figure 6.**
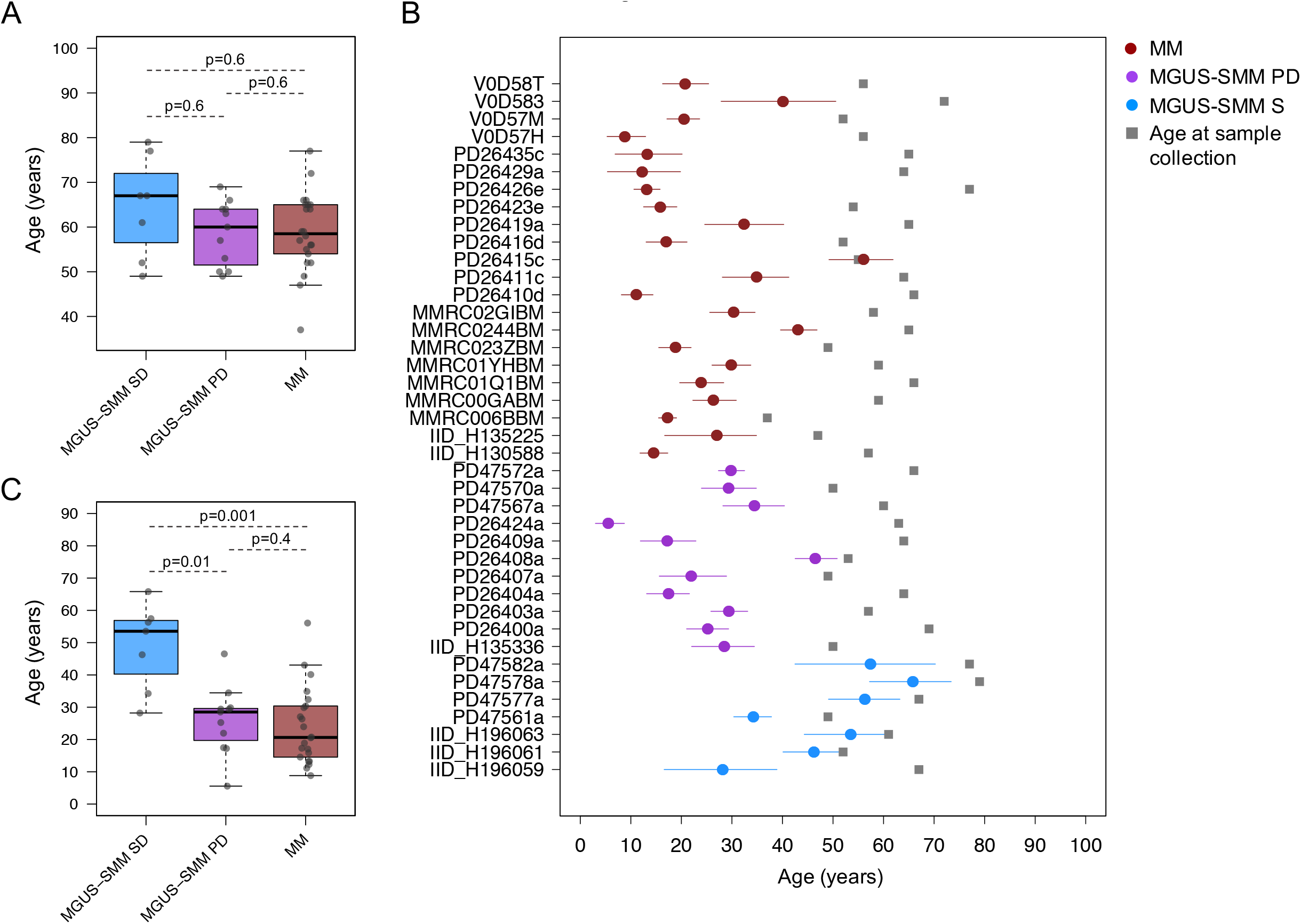
Timing the acquisition of the first multi-gain event in MM and myeloma precursor conditions. A) Comparison of the patients’ age at the time of sample collection between stable, progressive myeloma precursors and MM. p values were calculated using the Wilcoxon rank-sum test. B) Estimated patient age at the first multi-gain events with 95% confidence of intervals. Blue, purple and brown dots and lines represent stable, progressive myeloma precursors and MM respectively. Grey boxes reflect the sample collection time. C) Comparison of estimated patient age at the first multi-gain events between stable, progressive myeloma precursors and MM. p values were calculated using the Wilcoxon rank-sum test.

## Discussion

Early discovery work focusing on monoclonal serum proteins by Dr. Waldenstrom, Dr. Kyle, and others led to the emergence of two major schools of thought. Dr. Waldenstrom proposed that there were patients who had monoclonal proteins without any symptoms or evidence of end-organ damage, representing a benign monoclonal gammopathy.^42-45^ Conversely, the alternate opinion was that some patients with asymptomatic monoclonal proteins nevertheless progressed over time to MM, and that it was important to not term the process entirely benign. Thus in 1978, Dr. Kyle introduced the terminology MGUS, which allowed the field to move forward and to acknowledge the uncertainties in clinical outcome.^39^ The word “undetermined” was used to reflect the fact that, at diagnosis, it was not possible to determine which patients would ultimately progress to MM.

Over time, clinical risk scores for myeloma precursors conditions were developed based on indirect measurements of disease burden including BMPC percentage and the quantity of serum monoclonal protein).^3,5-7,9^ While such prognostic models have proven their utility, they have not been useful for identifying cases with MGUS and low-, and intermediate-risk SMM who may have already undergone malignant transformation.^5-7,9^

The historical differentiation between SMM and MGUS has been based on an arbitrary cut-point of 10% BMPC defined by immunohistochemistry. However, based on clinical experience, it is clear that some MGUS patients can progress rapidly despite their apparent low disease burden, and conversely many SMM patients will remain stable despite a higher disease burden with a behavior pattern typical of MGUS.^2,3,40,46^ An ability to recognize these two distinct clinical patterns independent of the BMPC percentage would offer significant advantages in clinical practice. The use of NGS has progressively provided an alternative to tumor burden-based models. Several studies have highlighted the importance of the value of genomic events for predicting progression of the myeloma precursor conditions. These studies have identified the value of mutations in the MAPK pathway and translocations in *MYC*.^5,13-15,18-20^ However, until recently, technical limitations (i.e. low number of clonal BMPC limiting the ability to conduct sequencing assays) led to most of these studies only including SMM cases and not MGUS. Here, thanks to the advent of multi-parametric BMPC flow-sorting and the application of low input WGS technology,^26,27^ we have been able to interrogate the WGS landscape of MGUS cases circumventing previous problems related to volume of clonal plasma cells and contamination by normal plasma cells. Given the ability of WGS to characterize SNV, SV, CNV and mutational signatures, we have shown that clinically stable cases of MGUS and SMM are characterized by a different genomic landscape and by differences in the temporal acquisition of myeloma-associated genomic events in comparison to progressive entities. The distribution of genetic events reveals the existence of two biologically and clinically distinct entities of asymptomatic monoclonal gammopathies: (i) one entity characterized by a sufficient number of myeloma genomic events to confer malignant potential and which is associated with progressive disease; and (ii) another entity with a lower burden of genetic events characterized by high likelihood of a prolonged, indolent, and clinically stable course.

Taken together, after more than 50 years of investigation of the relationship between myeloma precursor conditions and MM, the use of whole genome analysis has provided initial exiting evidence that myeloma precursor conditions with low disease burden at a high-risk of progression can be identified. Despite its limited sample size, this study provides proof of principle that WGS has the potential to accurately differentiate stable and progressive precursor conditions in low disease burden clinical states. The application of this technology in the clinic has the potential to significantly alter the management of individual patients but will require confirmation in larger studies. Going forward, improved and biology-oriented strategies to accurately identify patients with progressive myeloma precursor condition before clonal expansion i) will allow earlier initiation of therapy before onset of end-organ damage to avoid severe clinical complications; ii) will prevent patients with precursor conditions from being overtreated.^5,9^

## Methods

### Samples and whole genome sequencing

This study involved the use of human samples. Samples and data were obtained and managed in accordance with the Declaration of Helsinki. The study was approved by the medical ethics committees of the Jessa Hospital and Hasselt University (Belgium), Memorial Sloan Kettering Cancer Center (US) and Wellcome Sanger Institute (UK). Overall 26 samples of myeloma precursor condition and MM were newly sequenced for this study. Of these, 17 samples from 15 patients (15 MGUS, 1 SMM and 1 MM) were collected at the Jessa Hospital (**Supplemental Table 1**). One MGUS case (PD47563) and the only SMM case (PD47580) also had a second sample collected after 2 years of stable disease and at MM progression, respectively. Biological material from these cases used in this publication was provided by the Clinical Biobank of the Jessa Hospital and University Biobank Limburg (UBiLim).^47^ For all samples, BMPCs were isolated from bone marrow aspirates and sorted on a BD FACSAria IIITM instrument (BD Biosciences, San Jose, CA) using the markers CD19, CD20, CD38, CD45, CD56, CD138, CyIgL (BD Biosciences) and CyIgK (Agilent Technologies, Santa Clara, CA). Importantly, gating on of the general BMPC population was followed by gating on the pathological light chain - kappa or lambda-, according to the monoclonal protein in serum. Finally, CD56 positive or negative cells were selected depending on the patient characteristics, yielding a pure population of immunophenotypically aberrant PCs for sorting. For matched control DNA from each patient, bone marrow T cells or peripheral blood mononuclear cells were used. The T cells were isolated from the BM aspirates and sorted using the BD FACSAria IIITM (BD Biosciences) with anti-CD4 antibodies (BD Biosciences). Overall, we collected a median number of 3000 clonal BMPC per patient (range 1490-6000; **Supplemental Table 2**). This number of cells was too low to perform a standard WGS approach, and to overcome this, we used the recently published low input DNA enzymatic fragmentation-based WGS, which has been shown to have high accuracy in defining the WGS landscape of normal tissues from few thousands cells (**Figure 1A, Supplemental Table 1** and **Supplemental Methods**).^26,27^

For the remaining 8 newly sequenced cases (4 MGUS, 2 SMM and 2 MM) we used leftover DNA extracted from CD138-positive BMPC previously collected for SNP array investigation routinely performed in diagnostic at MSKCC. Having adequate DNA amount (>200 ng) and clonal purity estimation from the previous cytogenetic characterization, these samples were sequenced using standard WGS approaches (**Supplemental Methods**). Plasma cell selection was performed by magnetic bead-selection from bone marrow. Peripheral blood mononuclear cells were used as matched control.

To further increase the sample size of our cohort, we included 89 published WGSs from 52 MM patients.^25,48,49^ For 11 patients, samples were collected both at the time of SMM and MM progression.^13^ Overall, in this study we investigated WGS data from a total of 32 patients with multiple myeloma precursor condition.

### Processing of whole genome sequencing data

Overall, the median sequence coverage was 38X (range 27-97X; **Supplemental Table 1**). Short insert paired-end reads/FASTQ files were aligned to the reference human genome (GRCh37) using Burrows–Wheeler Aligner, BWA (v0.5.9). All samples were uniformly analyzed by the whole-genome analysis bioinformatic tools developed at the Wellcome Sanger Institute. Specifically: CaVEMan was used for SNVs, indels were analyzed with a modified Pindel version 2.0, for the identification of CNVs, ASCAT (v2.1.1) and Battenberg were performed. To determine the tumor clonal architecture, and to model clusters of clonal and subclonal point mutations, the Dirichlet process (DP) was applied ^13,25,29^. BRASS was used to detect SVs through discordantly mapping paired-end reads (large inversions and deletions, translocations, and internal tandem duplication). Complex events such as chromothripsis, chromoplexy, templated insertions were defined as previously described.^22,24,36,37,50^ All SVs not part of a complex event were define as single.^22,24^

The list of myeloma driver genes (n=80) was generated merging the two largest driver discovery studies.^12,22^ The list of SV hotspots was created by adding *MAFB* to the catalogue of 68 SV hotspots recently identified by our group.^21,24^

### Mutational signature analysis

Analysis of SBS signatures was performed following three main steps: 1) *de novo* extraction; 2) assignment; and 3) fitting.^29^ For the *de novo* extraction of mutational signatures we ran two independent algorithms; SigProfiler and the hierarchical Dirichlet process (hdp) (**Supplemental Figure 1**).^25,28^ Next, each extracted process was assigned to one or more mutational signatures included in the latest COSMIC v3.1 catalog (https://cancer.sanger.ac.uk/cosmic/signatures/SBS/index.tt). Lastly, *mmsig*, a fitting algorithm designed for hematological cancers (https://github.com/evenrus/mmsig), was applied to accurately estimate the contribution of each mutational signature in each sample.

### Molecular time

The relative timing of each multi-chromosomal gain event was estimated using the R package *mol_time* (https://github.com/nicos-angelopoulos/mol_time).^22^ This approach allows the estimation of the relative timing of acquisition of all large chromosomal gains (e.g. trisomies in hyperdiploid myeloma patients) using the corrected ratio between duplicated mutations (variant allele frequency; VAF 66%, acquired before the chromosomal duplication) and non-duplicated mutations (VAF 33%, acquired on either the non-duplicated allele or on one of the two duplicated alleles). Each clonal mutation VAF was corrected for the cancer purity estimated combining SNV and CNV data (**Supplementary Table 1**). Only chromosomal segments larger than 1 Mb and with more than 50 clonal mutations as estimated by the DP were considered.^22,25^ Tetrasomies, with both alleles duplicated, were removed given the impossibility of defining whether the two chromosomal gains occurred in close temporal succession or not.^22,25^ Using this approach, we were able to define if different chromosomal gains were acquired in the same or different time windows. Next, to convert the relative molecular time estimate into an absolute estimate, we combined chromosomal gains acquired in the same time window and re-calculated the molecular time using only the mutational burden of SBS1 and SBS5. These mutational processes are known to act in a constant way over time (i.e. clock-like) in MM (as in all cancers and normal tissues),^28,51^ and due to this feature we can convert the SBS1 and SBS5-based molecular clock into an absolute time estimate for the acquisition of these events in each patient’s life.^25,38,39^ Confidence of intervals were generated by bootstrapping 1000 times the molecular time estimate. Only multi-gain events with more than 50 SBS1 and SBS5 clonal mutations were included.

### Validation cohorts

To expand our CNV investigations, we included a validation cohort of 66 stable myeloma precursor condition, 2 progressive myeloma precursor condition, and 148 MM patients, with available SNP array data at the MSKCC (**Supplemental Table 7-8**). All cytogenetic data were reanalyzed using ASCAT (https://github.com/Crick-CancerGenomics/ascat).

To expand our investigations on nonsynonymous mutations and mutations in MM driver genes, WXS data from 33 MGUS patients were imported from EGA (EGAS00001001658)^19^ and analyzed using Caveman for SNVs and Pindel for indels. The copy number profile of each case was reconstructed using Facets. Finally, we also included as additional validation set 947 newly diagnosed MM enrolled in CoMMpass trial (AI15; NCT01454297; phs000748.v1.p1). The CoMMpass data were generated as part of the Multiple Myeloma Research Foundation Personalized Medicine Initiative (https://research.themmrf.org).

### Data analysis and statistics

Data analysis was carried out in R version 3.6.1. Standard statistical tests are mentioned consecutively in the manuscript while more complex analyses are described above. Wilcoxon rank-sum test between three groups was run using *pairwise*.*wilcox*.*test* R function with all p value adjusted for FDR. All reported p-values are two-sided, with a significance threshold of < 0.05.

## Data availability

Sequence files are available at the European Genome-phenome and dbGaP archive under the Accession codes:

- EGAD00001003309: 67 WGS data from 30 multiple myeloma patients
- phs000748.v1.p1: WXS and low coverage/long insert WGS sequencing data from 746 newly diagnosed multiple myeloma patients included in this study (CoMMpass trial; IA 15)
- phs000348.v2.p1 WGS data from 22 multiple myeloma patients
- EGAS00001001658: 33 WXS MGUS data

## Acknowledgements

We want to thank Dr. Vincent S Rajkumar for important and useful scientific feedback on the final draft.

This work is supported by the Memorial Sloan Kettering Cancer Center NCI Core Grant (P30 CA 008748), the Multiple Myeloma Research Foundation (MMRF), and the Perelman Family Foundation.

F.M. is supported by the American Society of Hematology, the International Myeloma Foundation and The Society of Memorial Sloan Kettering Cancer Center.

N.B. is funded by the European Research Council under the European Union’s Horizon 2020 research and innovation program (grant agreement no. 817997).

K.H.M. is supported by the Royal Australasian College of Physicians Dr Helen Rarity McCreanor Travelling Fellowship.

This study is part of the Limburg Clinical Research Center (LCRC) UHasselt-ZOL-Jessa, supported by the foundation Limburg Sterk Merk (LSM), Hasselt University, Ziekenhuis Oost-Limburg and Jessa Hospital. Additionally, this research is supported by Limburgs Kankerfonds, and LiveALife.

## Author contributions

F.M. designed and supervised the study, collected and analyzed the data and wrote the paper. G.F., N.B., J.L.R. and O.L. designed and supervised the study, collected the data and wrote the paper. B.O. designed the study, collected and analyzed the data and wrote the paper. K.H.M., D.L., P.C, V.Y., F.A., A.D., B.T.D. analyzed the data. B.Z.L., E.G., I.A., B.M., K.V., M.H., E.E.M., D.K., A.D., Y.Z., A.M., B.W. and G.M. collected the data. All authors read, revised, proved the manuscript

## References

1. Landgren, O. & Weiss, B.M. Patterns of monoclonal gammopathy of undetermined significance and multiple myeloma in various ethnic/racial groups: support for genetic factors in pathogenesis. Leukemia 23, 1691–1697 (2009).

2. Kyle, R.A., et al. Long-Term Follow-up of Monoclonal Gammopathy of Undetermined Significance. N Engl J Med 378, 241–249 (2018).

3. Kyle, R.A., et al. Clinical course and prognosis of smoldering (asymptomatic) multiple myeloma. N Engl J Med 356, 2582–2590 (2007).

4. Kyle, R.A., et al. Prevalence of monoclonal gammopathy of undetermined significance. N Engl J Med 354, 1362–1369 (2006).

5. Maura, F., et al. Moving From Cancer Burden to Cancer Genomics for Smoldering Myeloma: A Review. JAMA Oncol (2019).

6. Rajkumar, S.V., Landgren, O. & Mateos, M.V. Smoldering multiple myeloma. Blood 125, 3069–3075 (2015).

7. Rajkumar, S.V., et al. International Myeloma Working Group updated criteria for the diagnosis of multiple myeloma. Lancet Oncol 15, e538–548 (2014).

8. Cherry, B.M., et al. Modeling progression risk for smoldering multiple myeloma: results from a prospective clinical study. Leuk Lymphoma 54, 2215–2218 (2013).

9. Maura, F., Landgren, O. & Morgan, G.J. Designing Evolutionary Based Interception Strategies to Block the Transition from Precursor Phases to Multiple Myeloma. Clin Cancer Res (2020).

10. Cosemans, C., et al. Prognostic Biomarkers in the Progression From MGUS to Multiple Myeloma: A Systematic Review. Clin Lymphoma Myeloma Leuk 18, 235–248 (2018).

11. Manier, S., et al. Genomic complexity of multiple myeloma and its clinical implications. Nat Rev Clin Oncol 14, 100–113 (2017).

12. Walker, B.A., et al. Identification of novel mutational drivers reveals oncogene dependencies in multiple myeloma. Blood 132, 587–597 (2018).

13. Bolli, N., et al. Genomic patterns of progression in smoldering multiple myeloma. Nat Commun 9, 3363 (2018).

14. Bustoros, M., et al. Genomic Profiling of Smoldering Multiple Myeloma Identifies Patients at a High Risk of Disease Progression. J Clin Oncol, JCO2000437 (2020).

15. Lopez-Corral, L., et al. The progression from MGUS to smoldering myeloma and eventually to multiple myeloma involves a clonal expansion of genetically abnormal plasma cells. Clin Cancer Res 17, 1692–1700 (2011).

16. Lopez-Corral, L., et al. SNP-based mapping arrays reveal high genomic complexity in monoclonal gammopathies, from MGUS to myeloma status. Leukemia 26, 2521–2529 (2012).

17. Mailankody, S., et al. Baseline mutational patterns and sustained MRD negativity in patients with high-risk smoldering myeloma. Blood Adv 1, 1911–1918 (2017).

18. Mikulasova, A., et al. Genomewide profiling of copy-number alteration in monoclonal gammopathy of undetermined significance. Eur J Haematol 97, 568–575 (2016).

19. Mikulasova, A., et al. The spectrum of somatic mutations in monoclonal gammopathy of undetermined significance indicates a less complex genomic landscape than that in multiple myeloma. Haematologica 102, 1617–1625 (2017).

20. Misund, K., et al. MYC dysregulation in the progression of multiple myeloma. Leukemia 34, 322–326 (2020).

21. Barwick, B.G., et al. Multiple myeloma immunoglobulin lambda translocations portend poor prognosis. Nat Commun 10, 1911 (2019).

22. Maura, F., et al. Genomic landscape and chronological reconstruction of driver events in multiple myeloma. Nat Commun 10, 3835 (2019).

23. Maura, F., Rustad, E.H., Boyle, E.M. & Morgan, G.J. Reconstructing the evolutionary history of multiple myeloma. Best Pract Res Clin Haematol 33, 101145 (2020).

24. Rustad, E., et al. Revealing the impact of recurrent and rare structural variants in multiple myeloma. Blood Cancer Discovery in press.

25. Rustad, E.H., et al. Timing the initiation of multiple myeloma. Nat Commun 11, 1917 (2020).

26. Lee-Six, H., et al. The landscape of somatic mutation in normal colorectal epithelial cells. Nature 574, 532–537 (2019).

27. Moore, L., et al. The mutational landscape of normal human endometrial epithelium. Nature 580, 640–646 (2020).

28. Alexandrov, L.B., et al. The repertoire of mutational signatures in human cancer. Nature 578, 94–101 (2020).

29. Maura, F., et al. A practical guide for mutational signature analysis in hematological malignancies. Nat Commun 10, 2969 (2019).

30. Chan, K., et al. An APOBEC3A hypermutation signature is distinguishable from the signature of background mutagenesis by APOBEC3B in human cancers. Nat Genet 47, 1067–1072 (2015).

31. Maura, F., et al. Biological and prognostic impact of APOBEC-induced mutations in the spectrum of plasma cell dyscrasias and multiple myeloma cell lines. Leukemia (2017).

32. Walker, B.A., et al. APOBEC family mutational signatures are associated with poor prognosis translocations in multiple myeloma. Nat Commun 6, 6997 (2015).

33. Maura, F., et al. Role of AID in the temporal pattern of acquisition of driver mutations in multiple myeloma. Leukemia (2019).

34. Martincorena, I., et al. Universal Patterns of Selection in Cancer and Somatic Tissues. Cell 171, 1029–1041 e1021 (2017).

35. Kasar, S., et al. Whole-genome sequencing reveals activation-induced cytidine deaminase signatures during indolent chronic lymphocytic leukaemia evolution. Nat Commun 6, 8866 (2015).

36. Korbel, J.O. & Campbell, P.J. Criteria for inference of chromothripsis in cancer genomes. Cell 152, 1226–1236 (2013).

37. Li, Y., et al. Patterns of somatic structural variation in human cancer genomes. Nature 578, 112–121 (2020).

38. Gerstung, M., et al. The evolutionary history of 2,658 cancers. Nature 578, 122–128 (2020).

39. Mitchell, T.J., et al. Timing the Landmark Events in the Evolution of Clear Cell Renal Cell Cancer: TRACERx Renal. Cell 173, 611–623 e617 (2018).

40. Landgren, O., et al. Association of Immune Marker Changes With Progression of Monoclonal Gammopathy of Undetermined Significance to Multiple Myeloma. JAMA Oncol (2019).

41. Murray, D., et al. Detection and prevalence of monoclonal gammopathy of undetermined significance: a study utilizing mass spectrometry-based monoclonal immunoglobulin rapid accurate mass measurement. Blood Cancer J 9, 102 (2019).

42. Heremans, J.F., et al. Studies on “abnormal” serum globulins (M-components) in myeloma, macroglobulinemia and related diseases. Acta Med Scand Suppl 367, 1–126 (1961).

43. Kyle, R.A. Monoclonal gammopathy of undetermined significance. Natural history in 241 cases. Am J Med 64, 814–826 (1978).

44. Kyle, R.A. & Greipp, P.R. Smoldering multiple myeloma. N Engl J Med 302, 1347–1349 (1980).

45. Waldenstrom, J.G. Benign monoclonal gammapathy. Acta Med Scand 216, 435–447 (1984).

46. Lakshman, A., et al. Risk stratification of smoldering multiple myeloma incorporating revised IMWG diagnostic criteria. Blood Cancer J 8, 59 (2018).

47. Linsen, L., et al. Raising to the Challenge: Building a Federated Biobank to Accelerate Translational Research-The University Biobank Limburg. Front Med (Lausanne) 6, 224 (2019).

48. Chapman, M.A., et al. Initial genome sequencing and analysis of multiple myeloma. Nature 471, 467–472 (2011).

49. Lohr, J.G., et al. Widespread genetic heterogeneity in multiple myeloma: implications for targeted therapy. Cancer Cell 25, 91–101 (2014).

50. Maciejowski, J., et al. APOBEC3-dependent kataegis and TREX1-driven chromothripsis during telomere crisis. Nat Genet (2020).

51. Alexandrov, L.B., et al. Clock-like mutational processes in human somatic cells. Nat Genet 47, 1402–1407 (2015).

