## Supplemental Methods for "Whole genome sequencing provides evidence of two biologically and clinically distinct entities of asymptomatic monoclonal gammopathies: progressive versus stable myeloma precursor condition"

#### Low input whole genome sequencing.

The sorted cells were thawed on ice and after counting, the cells (range: 1,490-6,000) were centrifuged for 5 minutes at 400g. Cells were washed with FACSflow and spinned again at 400g for 5 min. The pelleted cells were lysed and DNA extraction was performed using the Arcturus® PicoPure® DNA Extraction Kit (Thermo Fisher Scientific, Waltham, MA), according to the manufacturer's instructions with minor modifications. Briefly, 155µL reconstitution buffer was added to one vial of proteinase K to obtain the Extraction Solution. Cell pellets were reconstituted in 20µL Extraction Solution and incubated at 65°C for 3 hours and 75°C for 30 minutes. After cooling down to room temperature (RT), samples were stored at -20°C until use. The on ice thawed lysates were manually processed using the low-input enzymatic fragmentation-based library preparation method of the Wellcome Sanger Institute (25, 26). Each 20µL lysate was mixed with 50µL TE Buffer (Ambion; 10 mM Tris- HCl, 1 mM EDTA) (Thermo Fisher Scientific, Invitrogen) and 50µL AMPure XP beads (Beckman Coulter, Brea, CA) at RT. After resuspending and vortexing, the lysate mixtures were incubated 5 minutes for binding reaction and 5 minutes for magnetic bead separation. Next, the genomic DNA (gDNA) was washed twice with 75% ethanol. After resuspending in 26µL TE buffer, the bead/gDNA slurry was used directly for DNA library construction. This protocol was based on the instructions of the NEBNext® Ultra™ II FS Kit (New England BioLabs, Ipswich, MA) for DNA Library Prep. Each sample was mixed with 7µL NEBNext Ultra II FS Reaction Buffer (New England BioLabs) and 2µL NEBNext Ultra II FS Enzyme Mix (New England BioLabs), and incubated for 12 minutes at 37°C and 30 minutes at 65°C to perform DNA fragmentation, end-repair and A-tailing. Next, this FS Reaction Mixture was incubated for 20 minutes at RT (~20°C) with a mixture of 30μL NEBNext Ultra II Ligation Master Mix (New England BioLabs), 1μL NEBNext Ligation Enhancer (New England BioLabs), 2.25μL nuclease-free water (Sigma-Aldrich, Saint Louis, MO) and 0.25μL TSQ Adapters (Integrated DNA Technologies (IDT), Coralville, IA Next, adapter-ligated libraries were purified by adding 65μL AMPure XP beads (Beckman Coulter) to the mixture. Following binding reaction, magnetic bead separation, and washing twice with 75% ethanol, the beads were eluted in nuclease-free water (Sigma-Aldrich). For amplification by PCR, 25μL eluted DNA library was mixed with 25μL KAPA HiFi HotStart Ready Mix (2x) (KAPA Biosystems, Wilmington, MA) and IDU tags (IDT), and incubated in the thermal cycler at 95°C for 5 minutes, then 12 cycles of 98°C for 30 seconds, 65°C for 30 seconds and 72°C for 2 minutes, and finally 72°C for 10 minutes. The amplified libraries were purified using 0.7:1 volumetric ratio of AMPure XP beads (Beckman Coulter) to PCR product. After the binding reaction, magnetic bead separation, and washing twice with 75% ethanol, the DNA libraries were eluted into nuclease-free water (Sigma-Aldrich) to obtain a final volume of 25μL (25, 26). Quantification and quality control of the DNA libraries was performed with Qubit™ dsDNA High Sensitivity Assay Kit (Thermo Fisher Scientific) on Qubit™ (Thermo Fisher Scientific), FlashGel® DNA System (Lonza, Basel, Switzerland), Agilent High Sensitivity DNA Kit Guide (Agilent Technologies) on 2100 Bio-Analyzer (Agilent Technologies). The genomic libraries were stored at -20°C until sequencing.

Finally, the DNA libraries were pooled and adjusted, flowcells were prepared, and sequencing clusters were generated. At the Wellcome Sanger Institute, the 150 base pairs (bp) paired-end sequencing was performed on the NovaSeq 150, S4 without XP (Illumina, San Diego, CA), according to the manufacturer's instructions

### Standard input whole genome sequencing

After PicoGreen quantification and quality control by Agilent BioAnalyzer, 500ng of genomic DNA were sheared using a LE220-plus Focused-ultrasonicator (Covaris catalog # 500569) and sequencing libraries were prepared using the KAPA Hyper Prep Kit (Kapa Biosystems KK8504) with modifications. Briefly, libraries were subjected to a 0.5X size select using aMPure XP beads (Beckman Coulter catalog # A63882) after post-ligation cleanup. Libraries not amplified by PCR (07652_C) were pooled equivolume and were quantitated based on their initial sequencing performance. Libraries amplified with 5 cycles of PCR (07652_D, 07652_F, 07652_G) were pooled equimolar. Samples were run on a NovaSeq 6000 in a 150bp/150bp paired end run, using the NovaSeq 6000 SBS v1 Kit and an S4 flow cell (Illumina).
