## Supplemental Figures for "Whole genome sequencing provides evidence of two biologically and clinically distinct entities of asymptomatic monoclonal gammopathies: progressive versus stable myeloma precursor condition"

Oben et al.

### Supplemental Figures

**Supplemental Figure 1.** De novo extraction of mutational signatures. The eight mutational signatures extracted by sigprofiler.

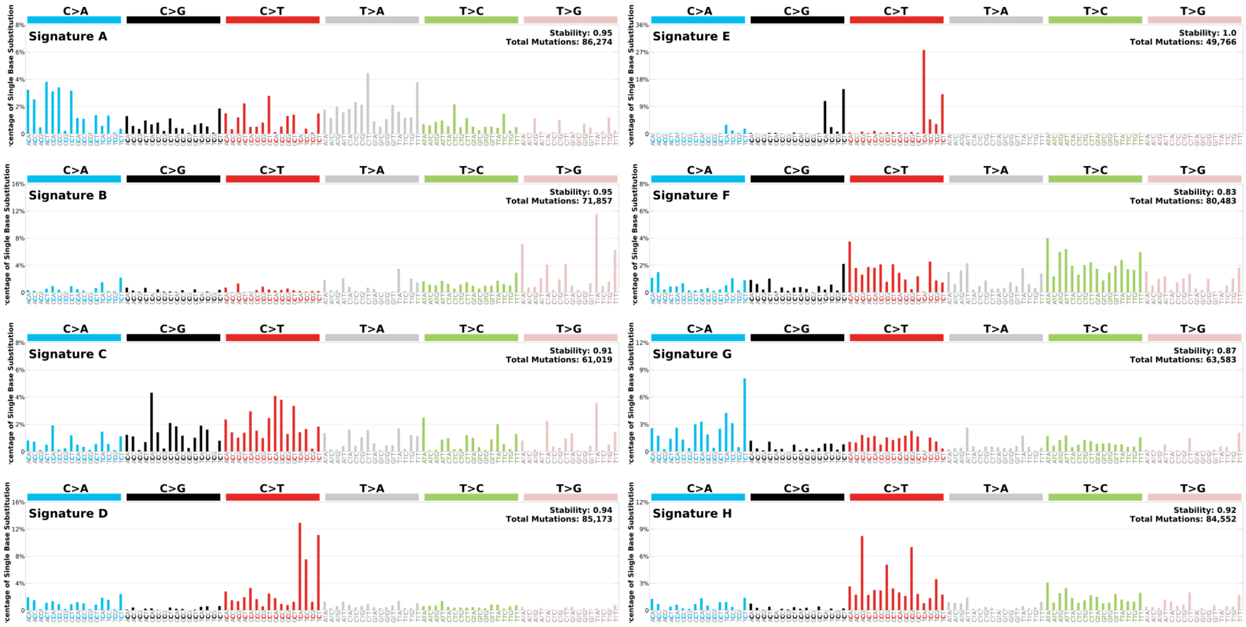

**Supplemental Figure 2.** The prevalence of recurrent multiple myeloma aneuploidies across the three groups investigated in this study: multiple myeloma (brown bars), stable and progressive myeloma precursor condition (blue and purple bars, respectively). Stable myeloma precursor condition showed a significantly lower prevalence of all recurrent cytogenetic aberration but 1p36 DEL (see **Supplemental Table 6**).

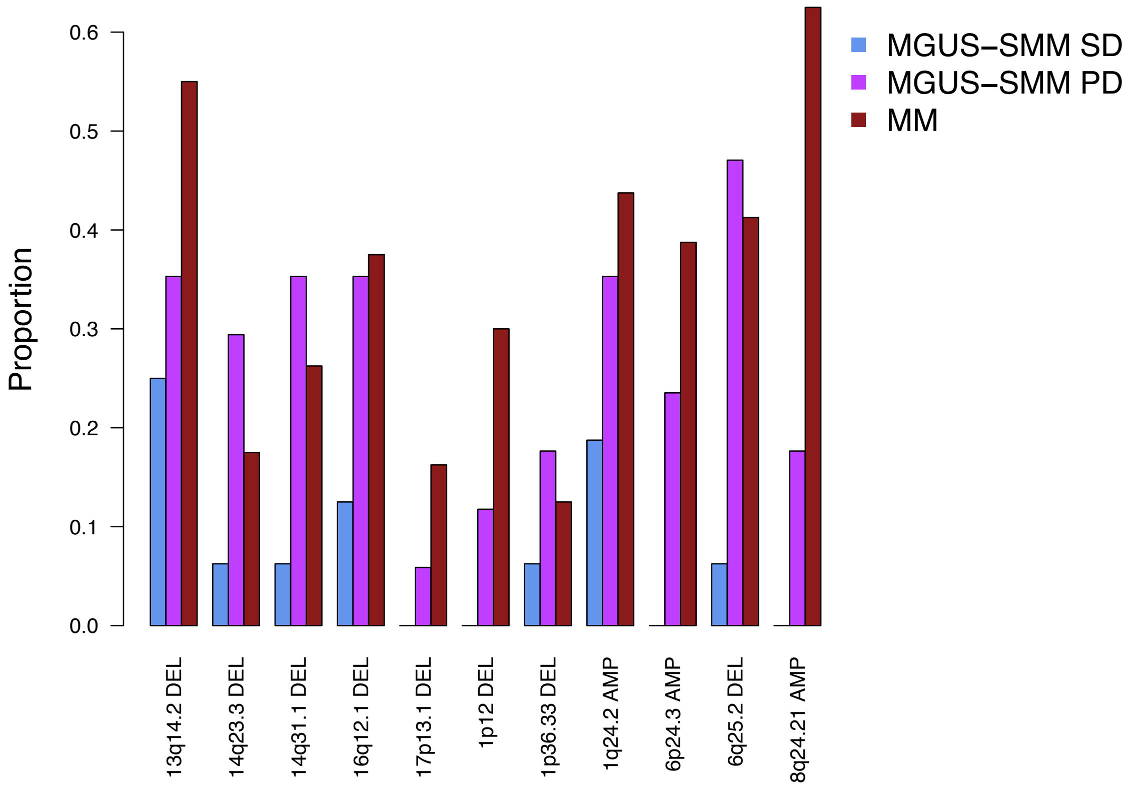

**Supplemental Figure 3.** Structural variants breakpoint distribution across the genome. Asterisks reflect significant association between SV and distinct genomic features tested using linear regression model (*lm* R function). Events involving *IGH* locus were excluded due to their known strong association with super enhancer.

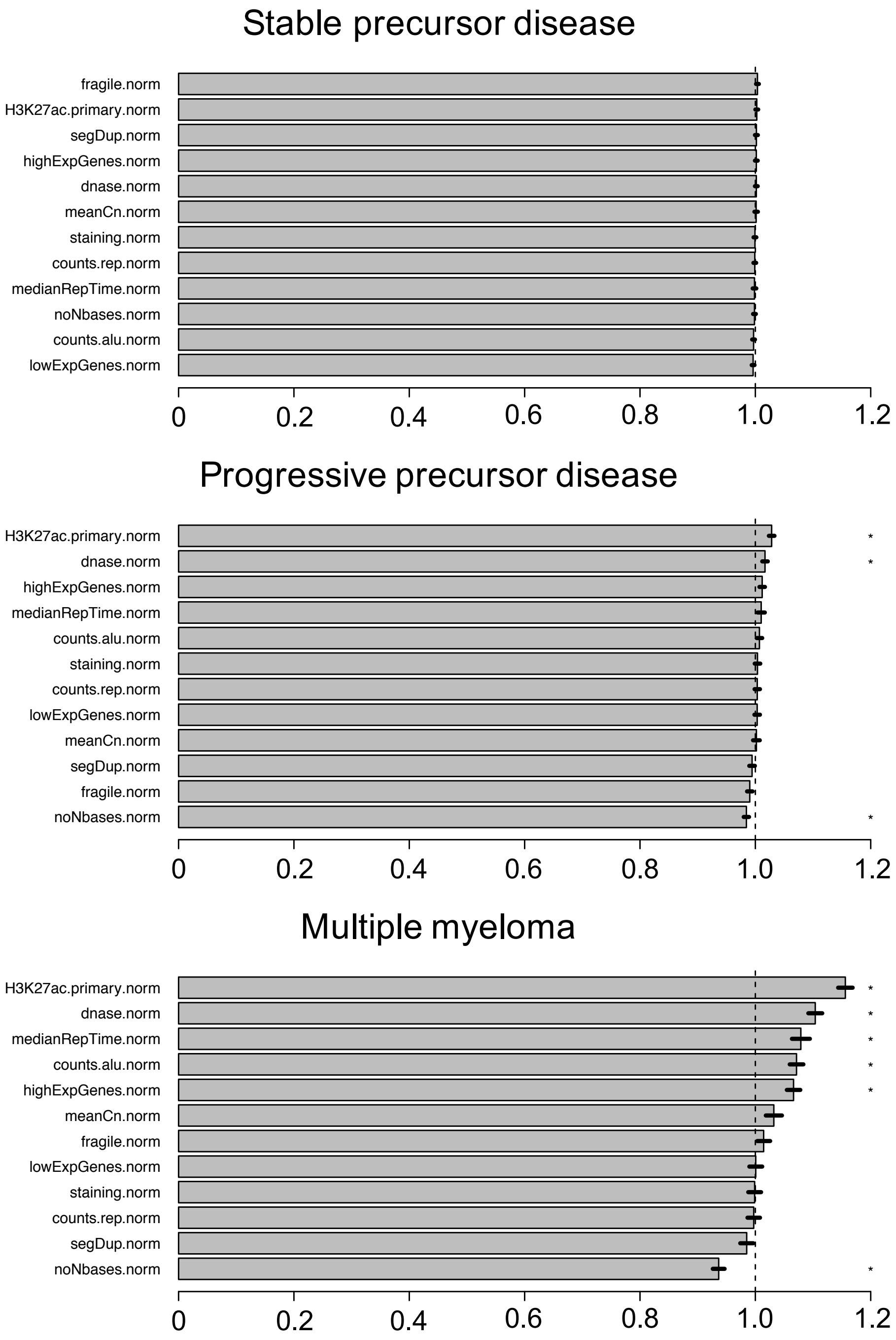

**Supplemental Figure 4.** Prevalence of known structural variants hotspots (n=69) across the three clinical stages: multiple myeloma (brown), stable and progressive myeloma precursor condition (blue and purple, respectively). p values were calculated using Wilcoxon rank-sum test

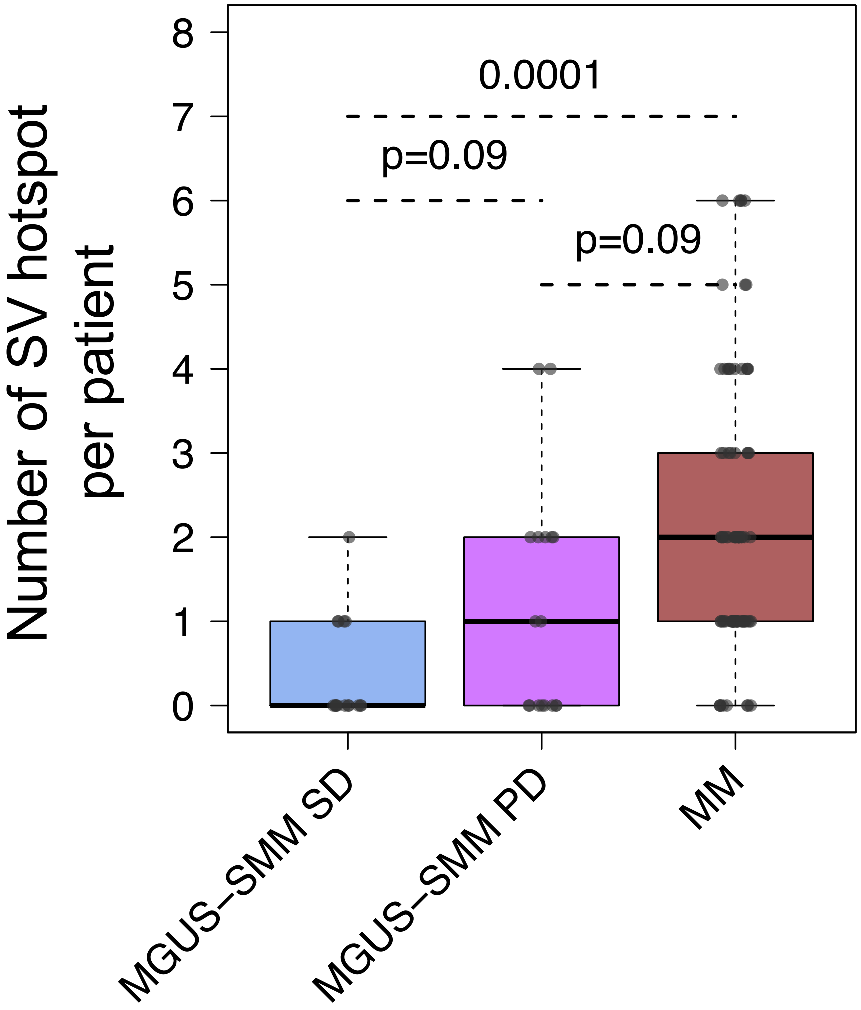

### Supplemental Tables

**Supplemental Table 2.** Summary of the total number of selected CD138+ bone marrow plasmacell sequenced using low input WGS approach.

| Sanger ID | Amount PCs | Amount cells matched control |
|---|---|---|
| **PD47575a** | 3000 | 6000 |
| **PD47582a** | 1518 | 2015 |
| **PD47561a** | 2285 | 6000 |
| **PD47577a** | 1000 | 3000 |
| **PD47578a** | 3000 | 3000 |
| **PD47579a** | 3000 | 3000 |
| **PD47563a** | 3000 | 3000 |
| **PD47563c** | 1495 | |
| **PD47573a** | 3000 | 3000 |
| **PD47581a** | 3000 | 3000 |
| **PD47574a** | 3000 | 6000 |
| **PD47576a** | 3000 | 3000 |
| **PD47567a** | 3000 | 3000 |
| **PD47570a** | 3000 | 3000 |
| **PD47572a** | 3000 | 3000 |
| **PD47580a** | 3000 | 3000 |
| **PD47580c** | 3000 | |

**Supplemental Table 3.** Percentage of bone marrow mononuclear cell at diagnosis for all cases with myeloma precursor condition included in this study. PD47576a has 10% BMPC at the first bone marrow evaluation, but <10% in all the subsequent follow up evaluations.

| Sample | Stage | BMPC (%) |
|---|---|---|
| **IID_H135336** | SMM - progressed | 10 |
| **IID_H196059** | MGUS - stable | 7 |
| **IID_H196061** | MGUS- stable | 4 |
| **IID_H196062** | SMM - stable | 15 |
| **IID_H196063** | MGUS- stable | 5 |
| **IID_H196064** | MGUS - progressed | 8 |
| **PD26400a** | SMM - progressed | 33 |
| **PD26401a** | SMM - progressed | 21 |
| **PD26402a** | SMM - progressed | 40 |
| **PD26403a** | SMM - progressed | 29 |
| **PD26404a** | SMM - progressed | 37 |
| **PD26405a** | SMM - progressed | 25 |
| **PD26406a** | SMM - progressed | NA |
| **PD26407a** | SMM - progressed | 39 |
| **PD26408a** | SMM - progressed | 31 |
| **PD26409a** | SMM - progressed | 10 |
| **PD26424a** | SMM - progressed | 61* |
| **PD47561a** | MGUS- stable | 3 |
| **PD47563a** | MGUS- stable | 3 |
| **PD47567a** | MGUS - progressed | 6.5 |
| **PD47570a** | MGUS - progressed | 5 |
| **PD47572a** | MGUS - progressed | 4.4 |
| **PD47573a** | MGUS- stable | 9.1 |
| **PD47574a** | MGUS- stable | 7 |
| **PD47575a** | MGUS- stable | 6 |
| **PD47576a** | MGUS- stable | 10 |
| **PD47577a** | MGUS- stable | 3.8 |
| **PD47578a** | MGUS- stable | 8.2 |
| **PD47579a** | MGUS- stable | 6 |
| **PD47580a** | SMM - progressed | 11.8 |
| **PD47581a** | MGUS- stable | 0.6 |
| **PD47582a** | MGUS- stable | 3.2 |

Key: MGUS: monoclonal gammopathy of undetermined significancep; SMM: smoldering multiple myeloma

*this case was diagnosed in 2010 prior to the 2014 IMWG diagnostic criteria for MM. Patients was monitored for high risk smoldering multiple myeloma with a less than 60% clonal plasmacells.

**Supplemental Table 4.** Assignment of mutational signatures extracted by *SigProfiler*.

| De novo extracted | Global NMF Signatures | Similarity |
|---|---|---|
| **Signature 96-A** | Signature SBS8 | 0.93 |
| **Signature 96-B** | Signature SBS9 | 0.93 |
| **Signature 96-C** | Signature SBS-MM1 | 1 |
| **Signature 96-D** | Signature SBS2 | 0.97 |
| **Signature 96-E** | Signature SBS2 (56.80%) & Signature SBS13 (43.20%) | 1 |
| **Signature 96-F** | Signature SBS5 | 0.93 |
| **Signature 96-G** | Signature SBS18 | 0.95 |
| **Signature 96-H** | Signature SBS1 (23.14%) & Signature SBS5 (76.86%) | 0.97 |

**Supplemental Table 5.** Catalogue of 80 driver genes mutated in multiple myeloma.

| Gene Symbol | Role | AID target |
|---|---|---|
| **ABCF1** | Unknown | |
| **ACTG1** | Unknown | |
| **ARID1A** | TSG | |
| **ARID2** | TSG | |
| **ATM** | TSG | |
| **ATRX** | TSG | |
| **BCL7A** | Oncogene | yes |
| **BHLHE41** | Unknown | |
| **BRAF** | Oncogene | |
| **BTG1** | TSG | yes |
| **CCND1** | Oncogene | |
| **CDKN1B** | TSG | |
| **CDKN2C** | TSG | |
| **CREBBP** | TSG | |
| **CYLD** | TSG | |
| **DIS3** | Oncogene | |
| **DNMT3A** | TSG | |
| **DTX1** | Unknown | yes |
| **DUSP2** | Unknown | yes |
| **EGR1** | Oncogene | yes |
| **EP300** | TSG | |
| **FAM46C** | TSG | |
| **FGFR3** | Oncogene | |
| **FUBP1** | TSG | |
| **HIST1H1B** | Unknown | yes |
| **HIST1H1D** | Unknown | yes |
| **HIST1H1E** | Oncogene | yes |
| **HIST1H2BK** | Unknown | yes |
| **HUWE1** | Unknown | |
| **IDH1** | Oncogene | |
| **IDH2** | Oncogene | |
| **IRF1** | Unknown | |
| **IRF4** | Oncogene | yes |
| **KDM5C** | TSG | |
| **KDM6A** | TSG | |
| **KLHL6** | Unknown | |
| **KMT2B** | TSG | |
| **KMT2C** | TSG | |
| **KRAS** | Oncogene | |
| **LCE1D** | Unknown | |

| | | |
|---|---|---|
| **LTB** | Unknown | yes |
| **MAF** | Oncogene | |
| **MAFB** | Oncogene | |
| **MAML2** | Oncogene | |
| **MAN2C1** | Unknown | |
| **MAX** | TSG | |
| **NCOR1** | TSG | |
| **NF1** | TSG | |
| **NFKB2** | Oncogene | |
| **NFKBIA** | TSG | |
| **NRAS** | Oncogene | |
| **PABPC1** | Unknown | |
| **PIK3CA** | Oncogene | |
| **PIM1** | Oncogene | yes |
| **POT1** | TSG | |
| **PRDM1** | TSG | |
| **PRKD2** | Unknown | |
| **PTPN11** | Oncogene | |
| **RASA2** | TSG | |
| **RB1** | TSG | |
| **RFTN1** | Unknown | |
| **RPL10** | TSG | |
| **RPL5** | TSG | |
| **RPRD1B** | Unknown | |
| **RPS3A** | Unknown | |
| **SAMHD1** | Unknown | |
| **SETD2** | TSG | |
| **SF3B1** | Oncogene | yes |
| **SP140** | TSG | |
| **TBC1D29** | Unknown | |
| **TCL1A** | Oncogene | |
| **TET2** | TSG | yes |
| **TGDS** | TSG | |
| **TP53** | TSG | |
| **TRAF2** | TSG | |
| **TRAF3** | TSG | |
| **UBR5** | TSG | |
| **XBP1** | Oncogene | yes |
| **ZFP36L1** | Unknown | yes |
| **ZNF292** | TSG | |

Key: TSG= tumor suppressor gene

**Supplemental Table 6.** Different prevalence of recurrent MM copy number changes across the three study groups: MM, stable and progressive MGUS-SMM. Fisher's test was used to estimate the p value.

| CNV event | p value<br>MGUS/SMM SD<br>vs<br>MGUS/SMM PD | p value<br>MGUS/SMM PD<br>vs<br>MM | p value<br>MGUS/SMM SD<br>vs<br>MM |
|---|---|---|---|
| **13q14.2 DEL** | 0.3908 | 0.1343 | <0.0001 |
| **14q23.3 DEL** | 0.0122 | 0.2063 | 0.0055 |
| **14q31.1 DEL** | 0.0028 | 0.4047 | <0.0001 |
| **16q12.1 DEL** | 0.0315 | 1 | <0.0001 |
| **17p13.1 DEL** | 0.1531 | 0.4849 | <0.0001 |
| **1p12 DEL** | 0.0222 | 0.1642 | <0.0001 |
| **1p36.33 DEL** | 0.1395 | 0.4637 | 0.13803 |
| **1q24.2 AMP** | 0.1965 | 0.616 | <0.0001 |
| **6p24.3 AMP** | 0.0003 | 0.3016 | <0.0001 |
| **6q25.2 DEL** | 0.0001 | 0.6218 | <0.0001 |
| **8q24.21 AMP** | 0.003 | 0.0004 | <0.0001 |

**Supplemental Table 7.** Summary of the clinical features of all cases with multiple myeloma with available SNP array data.

| Variable | Value |
|---|---|
| **Patients (n)** | 148 |
| **Age at diagnosis (years, median, IQR)** | 60.9 (54.3 - 69.1) |
| **Gender (female, male)** | 56, 92 |
| **ISS stage 1 (n, %)** | 30 (20.3%) |
| **2 (n, %)** | 60 (40.5%) |
| **3 (n, %)** | 30 (20.3%) |
| **NA (n, %)** | 28 (18.9%) |
| **Untreated (n, %)** | 87 (58.8%) |
| **Post-treatment (n, %)** | 61 (41.2%) |
| **PFS follow-up (years, median, IQR)** | 2.2 (1.5 - 3.0) |
| **Deaths (n, %)** | 20 (14%) |
| **OS follow-up (years, median, IQR)** | 3.1 (2.2 - 5.1) |

Key: ISS; International Staging System, IQR; inter-quartile range, NA; not available, OS; overall survival, PFS; progression-free survival

**Supplemental Table 8.** Summary of the clinical features of all cases with myeloma precursor condition with available SNP array data.

| **Variable** | **Value** |
|---|---|
| **Patients (n)** | 68 |
| **Age (median, IQR)** | 61.6 (53.6 - 67.9) |
| **Gender (female, male)** | 34, 34 |
| **MGUS (n, %)** | 15 (22.1) |
| **SMM (n, %)** | 53 (77.9) |
| **Plasma cells, >10% (n, %)** | 45 (66.2) |
| **>20% (n, %)** | 11 (16.2) |
| **M-spike, >2g/dL (n, %)** | 7 (10.3) |
| **>3g/dL (n, %)** | 2 (3.0) |
| **FLC ratio, >8 (n, %)** | 26 (38.2) |
| **>20 (n, %)** | 12 (17.6) |
| **HRCG (n, %)** | 9 (13.2) |
| **Non-IgG isotype (n, %)** | 25 (36.8) |
| **PFS follow-up (years, median, IQR)** | 2.1 (1.1 - 4.0) |

Key: HRCG; high risk cytogenetics (t(4;14), gain1q, del17p), FLC; free light chain, IQR; inter-quartile range, MGUS; monoclonal gammopathy of uncertain significance, SMM; smoldering multiple myeloma, PFS; progression-free survival
